# Chromosome-level genome assembly of Greenland scurvy-grass (*Cochlearia groenlandica*) reveals extensive centromeric satellite arrays

**DOI:** 10.64898/2026.11.05.764761

**Authors:** Ole K. Tørresen, Anne Krag Brysting, Bram Danneels, Rhianah Sandean, Spyridon Kollias, Giada Ferrari, Ave Tooming-Klunderud, Levi Yant, Kjetill S. Jakobsen

## Abstract

We present a chromosome-level, haplotype-resolved genome assembly from Greenland scurvy-grass (*Cochlearia groenlandica*). The assembly comprises two pseudo-haplotypes of 322 Mb and 295 Mb, with 89.7% and 95.6% respectively, each scaffolded into 7 chromosomes. Assembly completeness is high (BUSCO 97.8% in each haplotype), with 34,213 and 29,684 annotated protein-coding genes for haplotypes one and two, respectively. The chromosomes contain extensive centromeric satellite arrays that cover up to half of some chromosomes. Their precise functional boundaries remain undetermined.

## Introduction

The plant genus *Cochlearia* L. (Brassicaceae), with the common name scurvy grass, is known for its traditional use as a remedy to cure scurvy because of its high C-vitamin (ascorbic acid) content (Buckland et al., 1991). More recently, it has been developed as a model system to study polyploidy and local adaptation (Bray et al., 2024; Hämälä et al., 2024; Wolf et al., 2021) as it displays a range of interesting cytogenetic and ecotypic characteristics, including tolerance to cold, salinity, extreme bedrock types and heavy metal-contaminated soil (Eisenschmid et al., 2023; Nawaz et al., 2017; Wolf et al., 2021).

The genus, comprising 19 accepted species (WFO, 2026), is distributed in coastal and upland (alpine) habitats throughout Europe as well as in the circumpolar region. The plants are found in a wide range of natural habitats, from exposed coastal cliffs, salt marshes, sand dunes and freshwater springs to high-arctic tundra, and in addition in human-induced habitats such as metal-contaminated sites and salted roadways (Brandrud et al., 2017; Fekete et al., 2018; M. A. Koch, 2012). Most *Cochlearia* species are polyploid and the habitat differentiation is to some degree related to ploidy, with diploids usually found in upland freshwater springs or in high-arctic tundra, autotetraploids along the coast and higher ploidies (up to octoploid) in extreme salt marshes or human-induced habitats (Bray et al., 2024; Wolf et al., 2021). It is notable also that even diploid *Cochlearia* carries clear signals of mesohexaploidy (Mandáková et al., 2017).

Two base chromosome series exist, x=6 and x=7 (Gill, 1971; Saunte, 2010). Within the x=6 series, several diploid species are found in Central Europe, the British Isles and along the coast of Iceland. Based on meiotic cytology, the widespread tetraploid *C. officinalis*, the common scurvy grass, has recently been confirmed to be of autopolyploid origin (Bray et al., 2024), which is also a possible scenario for other, less widespread, tetraploids in the group. The evolutionary history of the higher ploidy levels is likely more complex and may involve both auto- and allopolyploidy (Hämälä et al., 2024; M. Koch, 2002; Wolf et al., 2021).

Within the x=7 group, the diploid *C. groenlandica* (Greenland scurvy-grass) is the most widespread species, with a circumpolar arctic distribution. It is mainly high-arctic but also found further south in subarctic regions. *Cochlearia* plants from across arctic North America, Russia-Siberia, Greenland and Svalbard all conform morphologically to *C. groenlandica* (Elven, 2011; Elven et al., 2020), and preliminary RAD sequencing data suggest a genetically coherent taxon, though with a clear geographical structure from Alaska in the west to Svalbard in the east (Bruholt, 2019). In addition to wet or dry, sandy or gravelly arctic tundra, *C. groenlandica* may also grow along seashores (Elven et al., 2020). In nutrient-rich bird-cliff meadows, it may be the dominant plant and grow much larger than in open tundra and other low-nutrient substrates (Eriksen & Nordal, 1989; Zmudczyńska-Skarbek et al., 2013). Several *Cochlearia* species are reported as self-incompatible, for instance tetraploid *C. officinalis* (Nordal & Stabbetorp, 1990) and hexaploid *C. bavarica* (Fischer et al., 2003; M. Koch, 2002). As opposed to this, *C. groenlandica* in Svalbard has been reported as self-compatible (Nordal & Laane, 1990), with correspondingly small flowers, lack of nectar and smell, as well as low insect visitation rate (Elven et al., 2020; Olsen et al., 2022).

Diploid (x=7) self-compatible plants that are genetically and morphologically close to *C. groenlandica* grow in alpine habitats in Iceland, where they are confined to late snow beds, ecologically similar to high-arctic tundra (Bruholt, 2019; Nordal & Laane, 1990; Olsen et al., 2022). Iceland is the only region where diploid plants of both base chromosome series coexist. In addition to the alpine plants, diploid plants of both chromosome numbers grow in coastal areas with beach cliffs. These plants are genetically and morphologically differentiated from the alpine plants (Olsen et al., 2022).

Here we present chromosome-scale, pseudo-haplotype-resolved assemblies of diploid *C. groenlandica*, representing the x=7 lineage of Cochlearia. Together with the previously published x=6 *C. excelsa* genome (Bray et al., 2024), this resource provides a foundation for investigating chromosome-number evolution, the complex origins of polyploid lineages, and genomic differentiation. It will also advance our possibility to untangle the evolutionary history of the higher ploidy species within the genus, as well as provide an improved basis for further studies on ecological adaptation.

## Material and Methods

### Sample acquisition and DNA extraction

Seeds, originally collected in 2009 by ALS Gustafsson from U.S.A., Alaska, Seward Peninsula, Wooly Lagoon N64° 51’ 18.7” W166° 24’ 6.7” (LG09-A-51; Gustafsson et al. (2022)), were germinated and the plant cultivated in a growth chamber at the University of Oslo. Approximately 2 g of leaf tissue were collected, immediately frozen and stored at -80 degree until DNA extraction.

Approximately 1 g of leaf tissue was ground with liquid nitrogen and washed with a 350 mM sorbitol buffer to osmotically remove secondary compounds (A. Jones et al., 2021), followed by CTAB lysis and DNA purification with Genomic-tip 100/G (QIAGEN). The complete protocol can be found at (T. L. Koch et al., 2026). Quality check of amount, purity and integrity of isolated DNA was performed using Qubit BR DNA quantification assay kit (Thermo Fisher), Nanodrop (Thermo Fisher), and Fragment Analyser (DNA HS 50kb large fragment kit, Agilent Tech.).

### Library preparation and sequencing for *de novo* assembly

Purified HMW DNA was sheared into an average fragment size of 15-20 kbp large fragments using the Megaruptor3 (Diagenode). A HiFi library was prepared following the PacBio protocol for HiFi library preparation using the SMRTbell® Prep Kit 3.0 and size-selected with a 10 kbp cut-off using a BluePippin (Sage Sciences). Sequencing was performed by the Norwegian Sequencing Centre on a PacBio Revio instrument (Pacific Biosciences Inc) with a 25M SMRT cell and Revio polymerase and sequencing kit, generating 16.9 Gb of HiFi data.

A Hi-C library was prepared by Dovetail Genomics LLC (Scotts Valley, California, USA) as part of an earlier Illumina short read version of the genome of the same plant (Grech et al., in prep). Whole plant tissue was flash frozen in liquid nitrogen and the Hi-C library was generated from whole tissue with endogenous chromatin extracted after fixation

### Genome assembly and curation, annotation and evaluation

A full list of relevant software tools and versions is presented in Table 1. We assembled the species using a pre-release of the EBP-Nor genome assembly pipeline (https://github.com/ebp-nor/GenomeAssembly). KMC (Kokot et al., 2017) was used to count k-mers of size 32 in the PacBio HiFi reads, excluding k-mers occurring more than 10,000 times. GenomeScope (Ranallo-Benavidez et al., 2020) was run as part of the pipeline on the k-mer histogram output from KMC and was included in the methods for completeness. Ploidy level was investigated using Smudgeplot (Ranallo-Benavidez et al., 2020). HiFiAdapterFilt (Sim et al., 2022) was applied on the HiFi reads to remove possible remnant PacBio adapter sequences. The filtered HiFi reads were assembled using Hifiasm (Cheng et al., 2021) with Hi-C integration resulting in a pair of haplotype-resolved assemblies, pseudo-haplotype one (hap1) and pseudo-haplotype two (hap2). Unique k-mers in each assembly/pseudo-haplotype were identified using meryl (Rhie et al., 2020) and used to create two sets of Hi-C reads, one without any k-mers occurring uniquely in hap1 and the other without k-mers occurring uniquely in hap2.

**Table 1.**
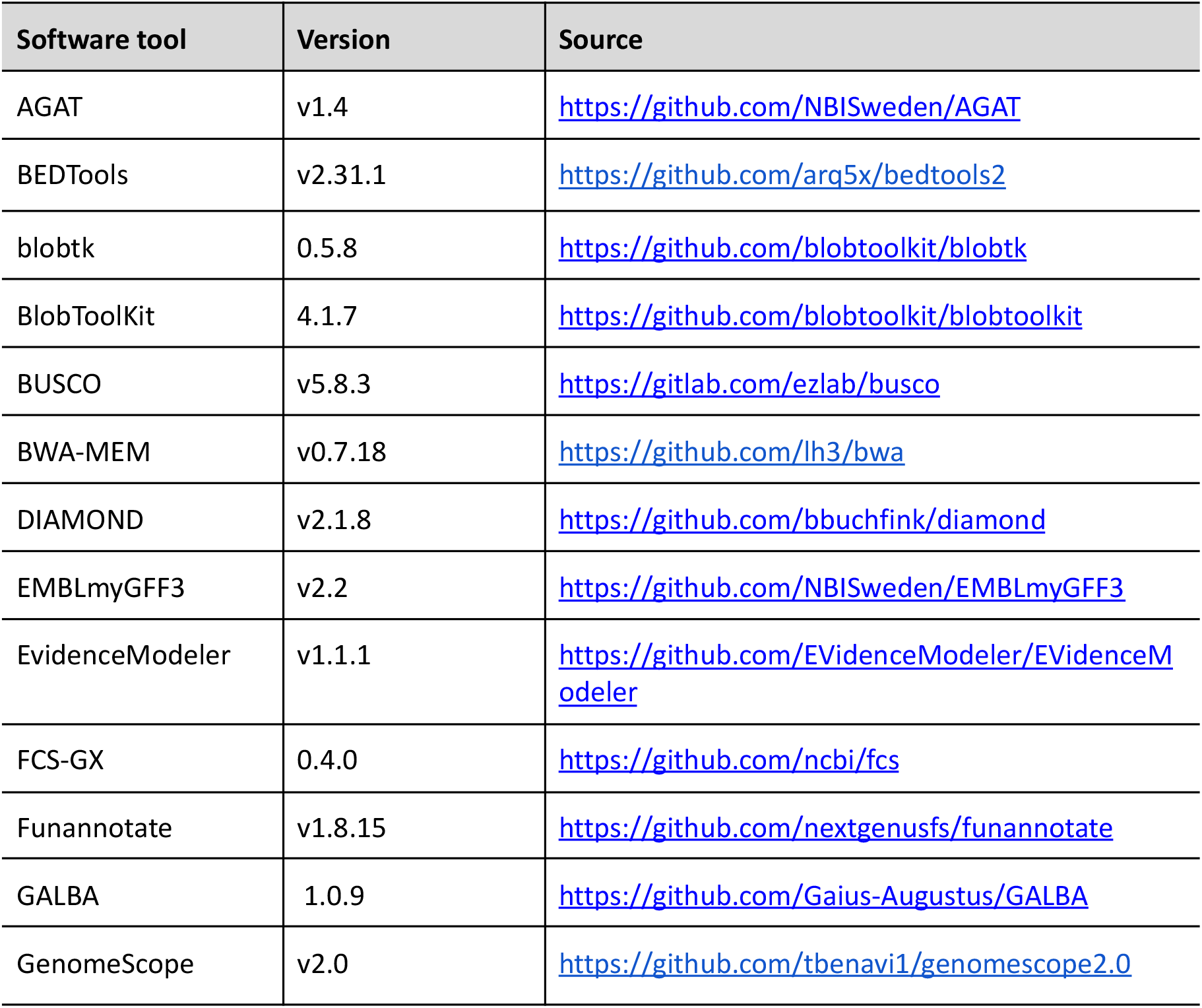
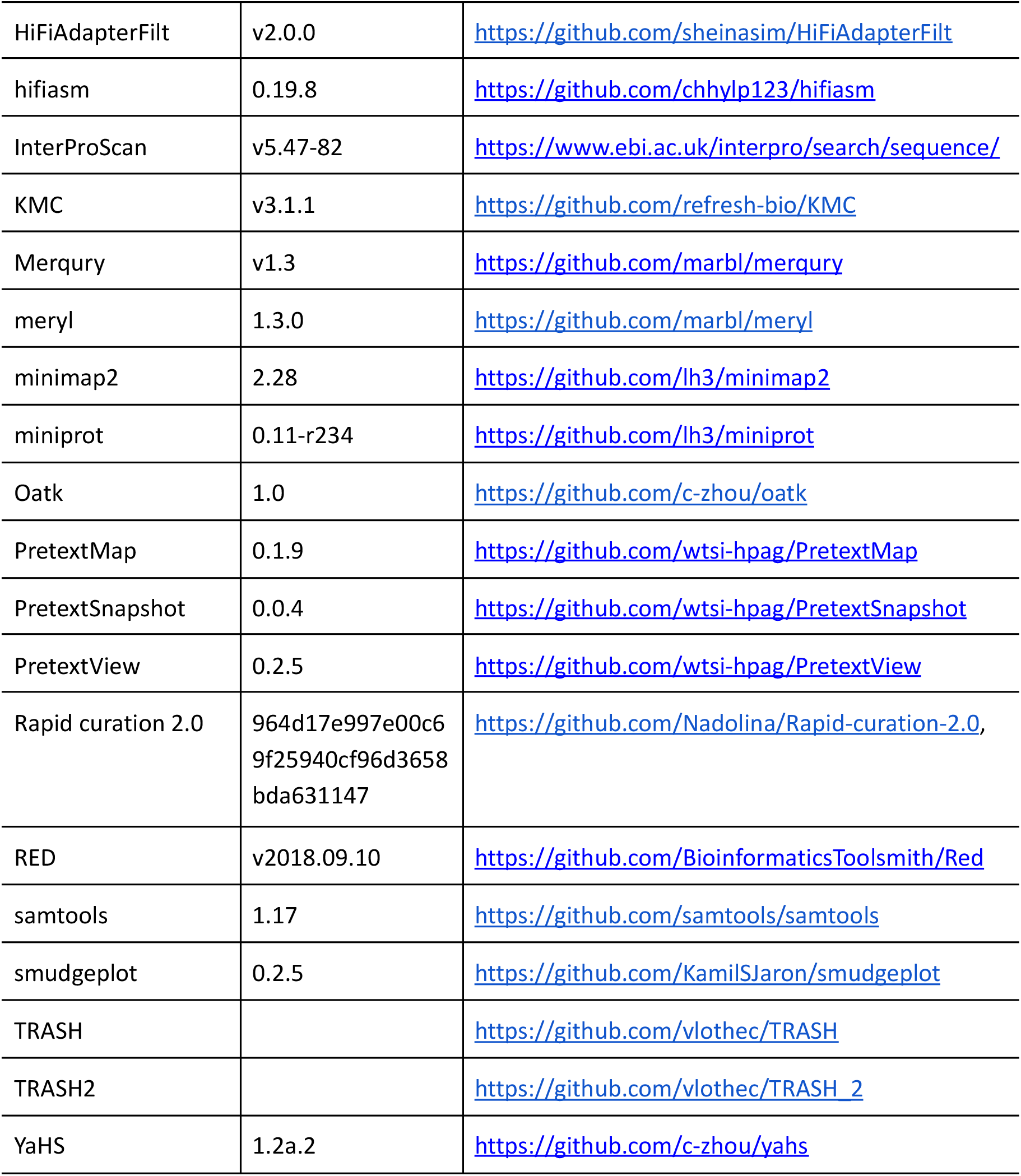
Software tools: versions and sources.

| Software tool | Version | Source |
|---|---|---|
| AGAT | v1.4 | https://github.com/NBISweden/AGAT |
| BEDTools | v2.31.1 | https://github.com/arq5x/bedtools2 |
| blobtk | 0.5.8 | https://github.com/blobtoolkit/blobtk |
| BlobToolKit | 4.1.7 | https://github.com/blobtoolkit/blobtoolkit |
| BUSCO | v5.8.3 | https://gitlab.com/ezlab/busco |
| BWA-MEM | v0.7.18 | https://github.com/lh3/bwa |
| DIAMOND | v2.1.8 | https://github.com/bbuchfink/diamond |
| EMBLmyGFF3 | v2.2 | https://github.com/NBISweden/EMBLmyGFF3 |
| EvidenceModeler | v1.1.1 | https://github.com/EVidenceModeler/EVidenceModeler |
| FCS-GX | 0.4.0 | https://github.com/ncbi/fcs |
| Funannotate | v1.8.15 | https://github.com/nextgenusfs/funannotate |
| GALBA | 1.0.9 | https://github.com/Gaius-Augustus/GALBA |
| GenomeScope | v2.0 | https://github.com/tbenavi1/genomescope2.0 |
| HiFiAdapterFilt | v2.0.0 | https://github.com/sheinasim/HiFiAdapterFilt |
| hifiasm | 0.19.8 | https://github.com/chhylp123/hifiasm |
| InterProScan | v5.47-82 | https://www.ebi.ac.uk/interpro/search/sequence/ |
| KMC | v3.1.1 | https://github.com/refresh-bio/KMC |
| Merqury | v1.3 | https://github.com/marbl/merqury |
| meryl | 1.3.0 | https://github.com/marbl/meryl |
| minimap2 | 2.28 | https://github.com/lh3/minimap2 |
| miniprot | 0.11-r234 | https://github.com/lh3/miniprot |
| Oatk | 1.0 | https://github.com/c-zhou/oatk |
| PretextMap | 0.1.9 | https://github.com/wtsi-hpag/PretextMap |
| PretextSnapshot | 0.0.4 | https://github.com/wtsi-hpag/PretextSnapshot |
| PretextView | 0.2.5 | https://github.com/wtsi-hpag/PretextView |
| Rapid curation 2.0 | 964d17e997e00c69f25940cf96d3658bda631147 | https://github.com/Nadolina/Rapid-curation-2.0, |
| RED | v2018.09.10 | https://github.com/BioinformaticsToolsmith/Red |
| samtools | 1.17 | https://github.com/samtools/samtools |
| smudgeplot | 0.2.5 | https://github.com/KamilSJaron/smudgeplot |
| TRASH |  | https://github.com/vlothec/TRASH |
| TRASH2 |  | https://github.com/vlothec/TRASH_2 |
| YaHS | 1.2a.2 | https://github.com/c-zhou/yahs |

K-mer filtered Hi-C reads were aligned to each scaffolded assembly using BWA-MEM (Li, 2013) with -5SPM options. The alignments were sorted based on name using samtools (Li et al., 2009) before applying samtools fixmate to remove unmapped reads and secondary alignments and to add mate score, and samtools markdup to remove duplicates. The resulting BAM files were used to scaffold the two assemblies using YaHS (Zhou et al., 2023) with default options. FCS-GX (Astashyn et al., 2024) was used to search for putative contamination and contaminated sequences were removed. The mitochondrion was searched for in contigs and reads using Oatk (Zhou et al., 2025). The assemblies were manually curated using PretextView and Rapid curation 2.0. Chromosomes were identified by inspecting the Hi-C contact map in PretextView and named according to size, with the longest as chromosome 1. We annotated the genome assemblies using a pre-release version of the EBP-Nor genome annotation pipeline (https://github.com/ebp-nor/GenomeAnnotation). First, AGAT (https://zenodo.org/record/7255559) `agat_sp_keep_longest_isoform.pl` and `agat_sp_extract_sequences.pl` were used on the *Arabidopsis thaliana* (TAIR10.1, GCF_000001735.4) genome assembly and annotation to generate one protein (the longest isoform) per gene. Miniprot (Li, 2023) was used to align the proteins to the curated assemblies. UniProtKB/Swiss-Prot (UniProt Consortium, 2023) release 2024_04 in addition to the Viridiplantae part of OrthoDB v11 (Kuznetsov et al., 2023) were also aligned separately to the assemblies. RED (Girgis, 2015) was run via redmask (https://github.com/nextgenusfs/redmask) on the assemblies to mask repetitive areas. GALBA (Brůna et al., 2023; Buchfink et al., 2015; Hoff & Stanke, 2019; Li, 2023; Stanke et al., 2006) was run with the *Arabidopsis* proteins using the miniprot mode on the masked assemblies.

The `funannotate-runEVM.py` script from Funannotate was used to run EvidenceModeler (Haas et al., 2008) on the alignments of GRCz11 proteins, UniProtKB/Swiss-Prot proteins, *Arabidposis* proteins and the predicted genes from GALBA. The resulting predicted proteins were compared to the protein repeats that Funannotate distributes using DIAMOND blastp and the predicted genes were filtered based on this comparison using AGAT. The filtered proteins were compared to the UniProtKB/Swiss-Prot release 2025_03 using DIAMOND (Buchfink et al., 2015) `blastp` to find gene names and InterProScan (P. Jones et al., 2014) was used to discover functional domains. AGATs `agat_sp_manage_functional_annotation.pl` was used to attach the gene names and functional annotations to the predicted genes. EMBLmyGFF3 (Norling et al., 2018) was used to combine the fasta files and GFF3 files into a EMBL format for submission to ENA.

All the evaluation tools have also been implemented in a pipeline, similar to assembly and annotation (https://github.com/ebp-nor/GenomeEvaluation). Merqury (Rhie et al., 2020) was used to assess the completeness and quality of the genome assemblies by comparing them to the k-mer content of both the Hi-C reads and PacBio HiFi reads. BUSCO (Manni et al., 2021) was used to assess the completeness of the genome assemblies by comparing against the expected gene content in the eudicots lineage. Gfastats (Formenti et al., 2022) was used to output different assembly statistics of the assemblies. BlobToolKit and BlobTools2 (Laetsch & Blaxter, 2017), in addition to blobtk were used to visualize assembly statistics. To generate the Hi-C contact map image, the Hi-C reads were mapped to the assemblies using BWA-MEM (Li, 2013) using the same approach as above. Finally, PretextMap was used to create a contact map which was visualized using PretextSnapshot.

Putative centromere sequences were identified with TRASH (Wlodzimierz et al., 2023) and TRASH2. Seaborn (Waskom, 2021) was used to visualise the satellite sequences together with exons by plotting the density of satellite sequences identified by TRASH2 together with the density of exons in 200 kb windows across the chromosomes.

Parts of the text in Methods and parts of Results is based on a template we use for all the species we publish in the EBP-Nor project.

## Results

### *De novo* genome assembly and annotation

The genome of *Cochlearia groenlandica* was assembled from a total of 57-fold coverage in Pacific Biosciences single-molecule HiFi long reads and 181-fold coverage in Arima Hi-C reads resulting in two haplotype-separated assemblies. The final assemblies have total lengths of 322 Mb and 295 Mb (Table 2 and Figure 2), respectively. Pseudo-haplotypes one (hap1) and two (hap2) have scaffold N50 size of 38.2 Mb and 46.9 Mb, respectively, and contig N50 of 7.8 Mb and 7.2 Mb, respectively (Table 2 and Fig. 2). Seven pseudo-chromosomes were identified in each pseudo-haplotype.

**Table 2:**
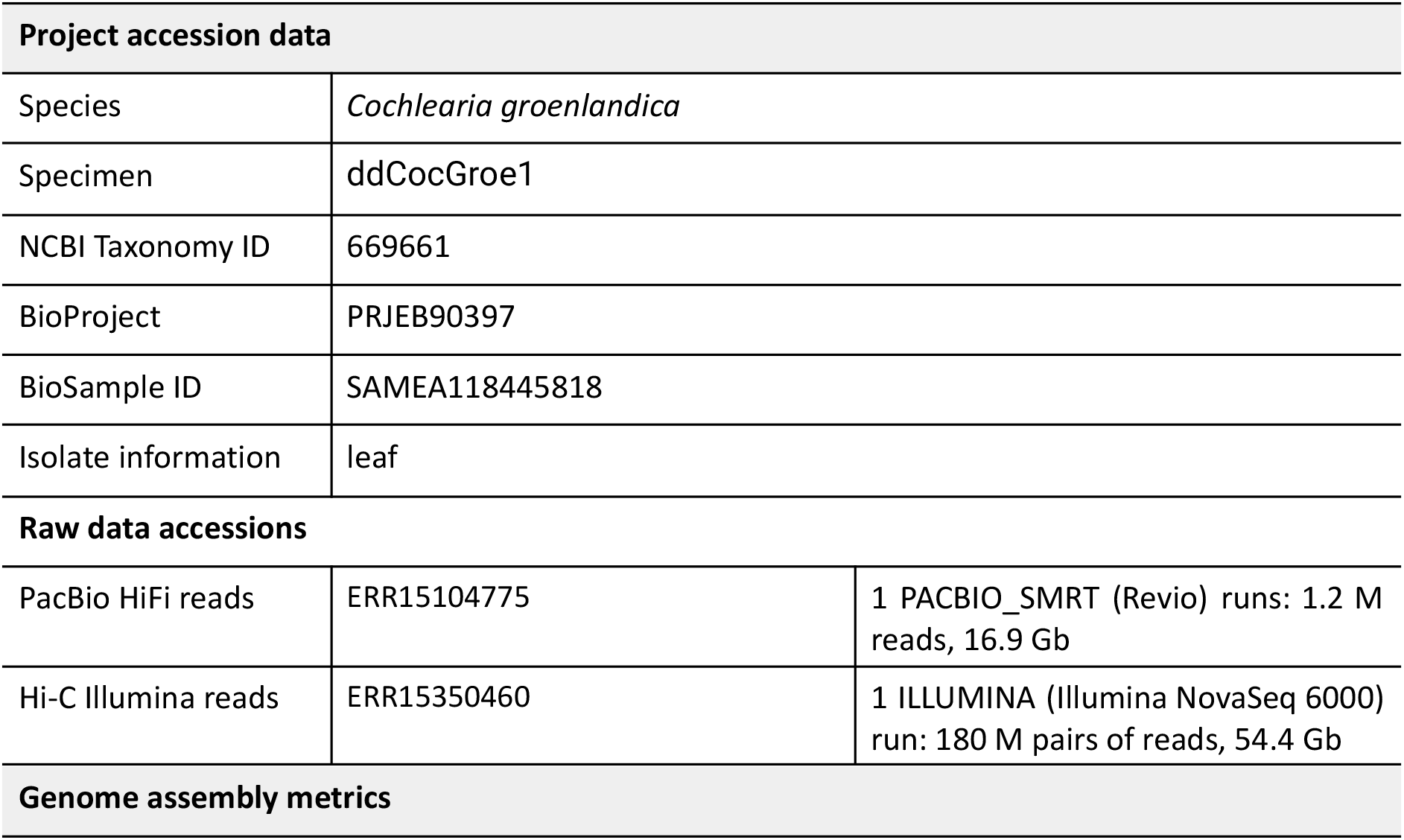
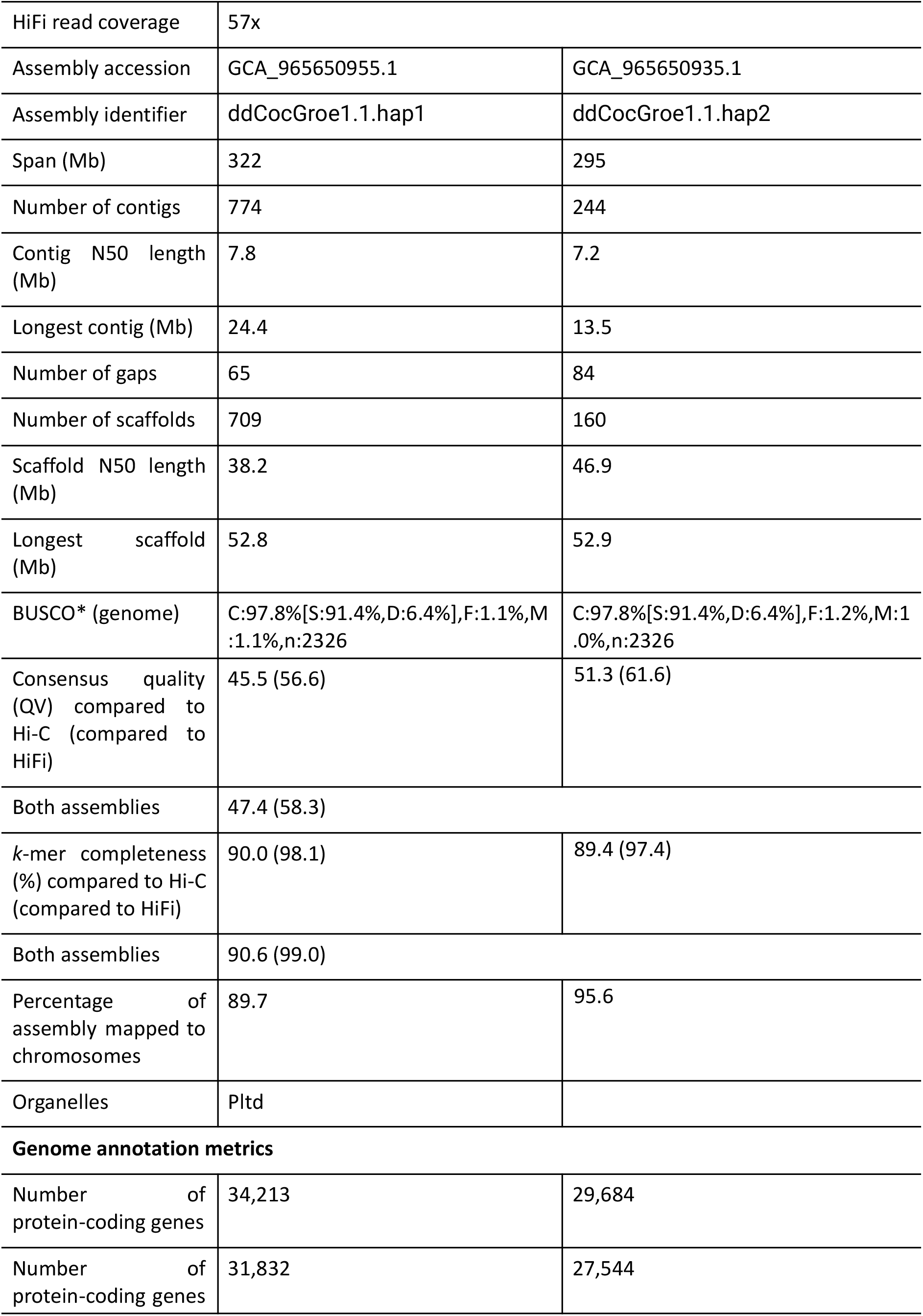
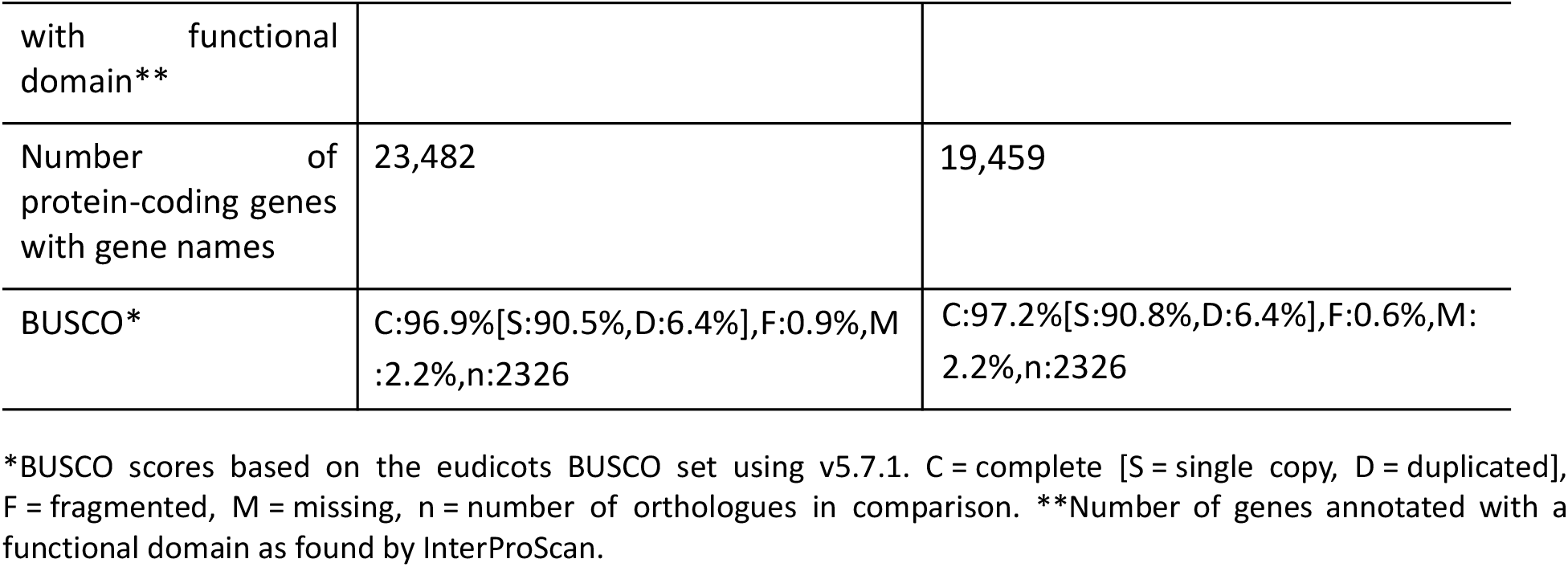
Genome data for *C. groenlandica*.

| **Project accession data** |  |  |
|---|---|---|
| Species | *Cochlearia groenlandica* |  |
| Specimen | ddCocGroe1 |  |
| NCBI Taxonomy ID | 669661 |  |
| BioProject | PRJEB90397 |  |
| BioSample ID | SAMEA118445818 |  |
| Isolate information | leaf |  |
| **Raw data accessions** |  |  |
| PacBio HiFi reads | ERR15104775 | 1 PACBIO_SMRT (Revio) runs: 1.2 M reads, 16.9 Gb |
| Hi-C Illumina reads | ERR15350460 | 1 ILLUMINA (Illumina NovaSeq 6000) run: 180 M pairs of reads, 54.4 Gb |
| **Genome assembly metrics** |  |  |
| HiFi read coverage | 57x |  |
| Assembly accession | GCA_965650955.1 | GCA_965650935.1 |
| Assembly identifier | ddCocGroe1.1.hap1 | ddCocGroe1.1.hap2 |
| Span (Mb) | 322 | 295 |
| Number of contigs | 774 | 244 |
| Contig N50 length (Mb) | 7.8 | 7.2 |
| Longest contig (Mb) | 24.4 | 13.5 |
| Number of gaps | 65 | 84 |
| Number of scaffolds | 709 | 160 |
| Scaffold N50 length (Mb) | 38.2 | 46.9 |
| Longest scaffold (Mb) | 52.8 | 52.9 |
| BUSCO* (genome) | C:97.8%[S:91.4%,D:6.4%],F:1.1%,M:1.1%,n:2326 | C:97.8%[S:91.4%,D:6.4%],F:1.2%,M:1.0%,n:2326 |
| Consensus quality (QV) compared to Hi-C (compared to HiFi) | 45.5 (56.6) | 51.3 (61.6) |
| Both assemblies | 47.4 (58.3) |  |
| *k*-mer completeness (%) compared to Hi-C (compared to HiFi) | 90.0 (98.1) | 89.4 (97.4) |
| Both assemblies | 90.6 (99.0) |  |
| Percentage of assembly mapped to chromosomes | 89.7 | 95.6 |
| Organelles | Pltd |  |
| **Genome annotation metrics** |  |  |
| Number of protein-coding genes | 34,213 | 29,684 |
| Number of protein-coding genes with functional domain** | 31,832 | 27,544 |
| Number of protein-coding genes with gene names | 23,482 | 19,459 |
| BUSCO* | C:96.9%[S:90.5%,D:6.4%],F:0.9%,M:2.2%,n:2326 | C:97.2%[S:90.8%,D:6.4%],F:0.6%,M:2.2%,n:2326 |

\*BUSCO scores based on the eudicots BUSCO set using v5.7.1. C = complete [S = single copy, D = duplicated], F = fragmented, M = missing, n = number of orthologues in comparison. \*\*Number of genes annotated with a functional domain as found by InterProScan.

**Figure 1:**
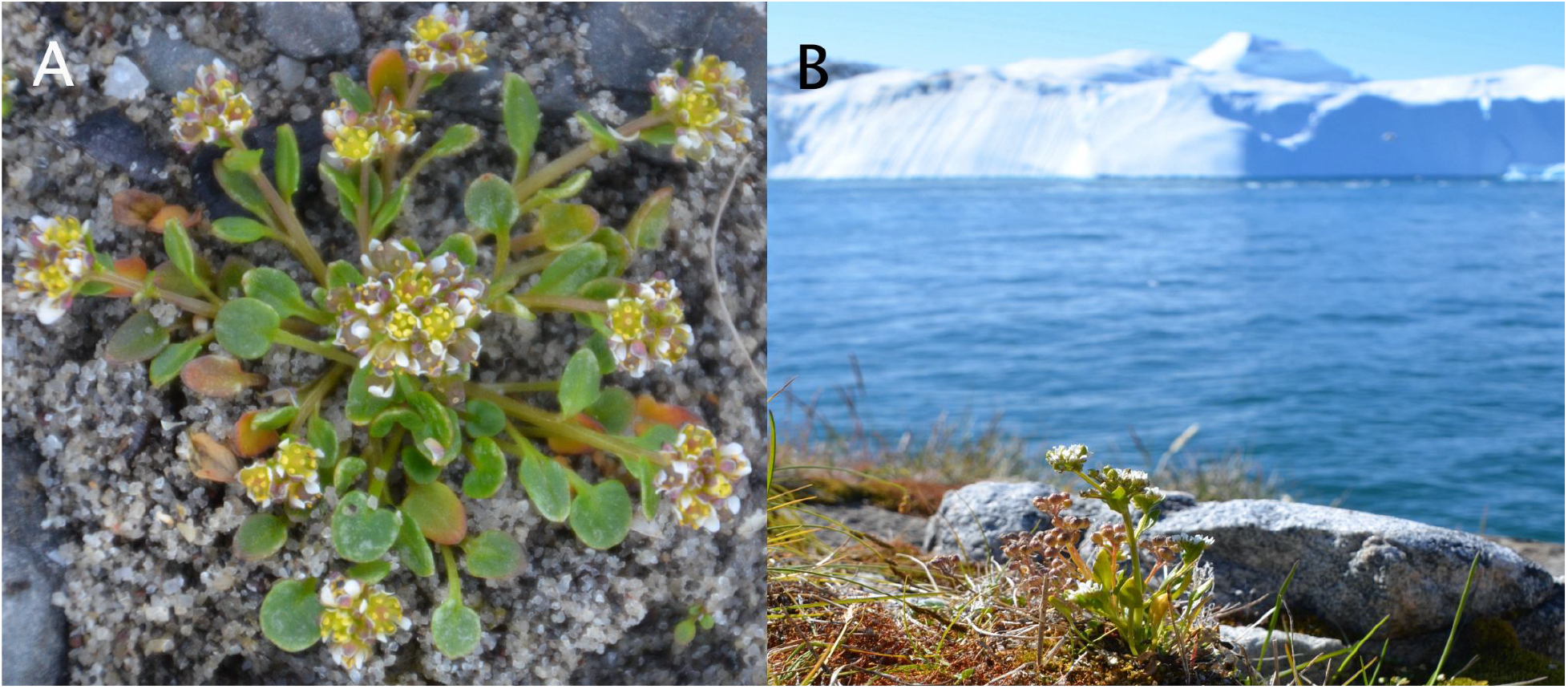
*Cochlearia groenlandica* in Western Greenland. A) A close-up showing the plant. B) The plant in its Arctic environment. Photo: Charlotte S. Bjorå.

**Figure 2:**
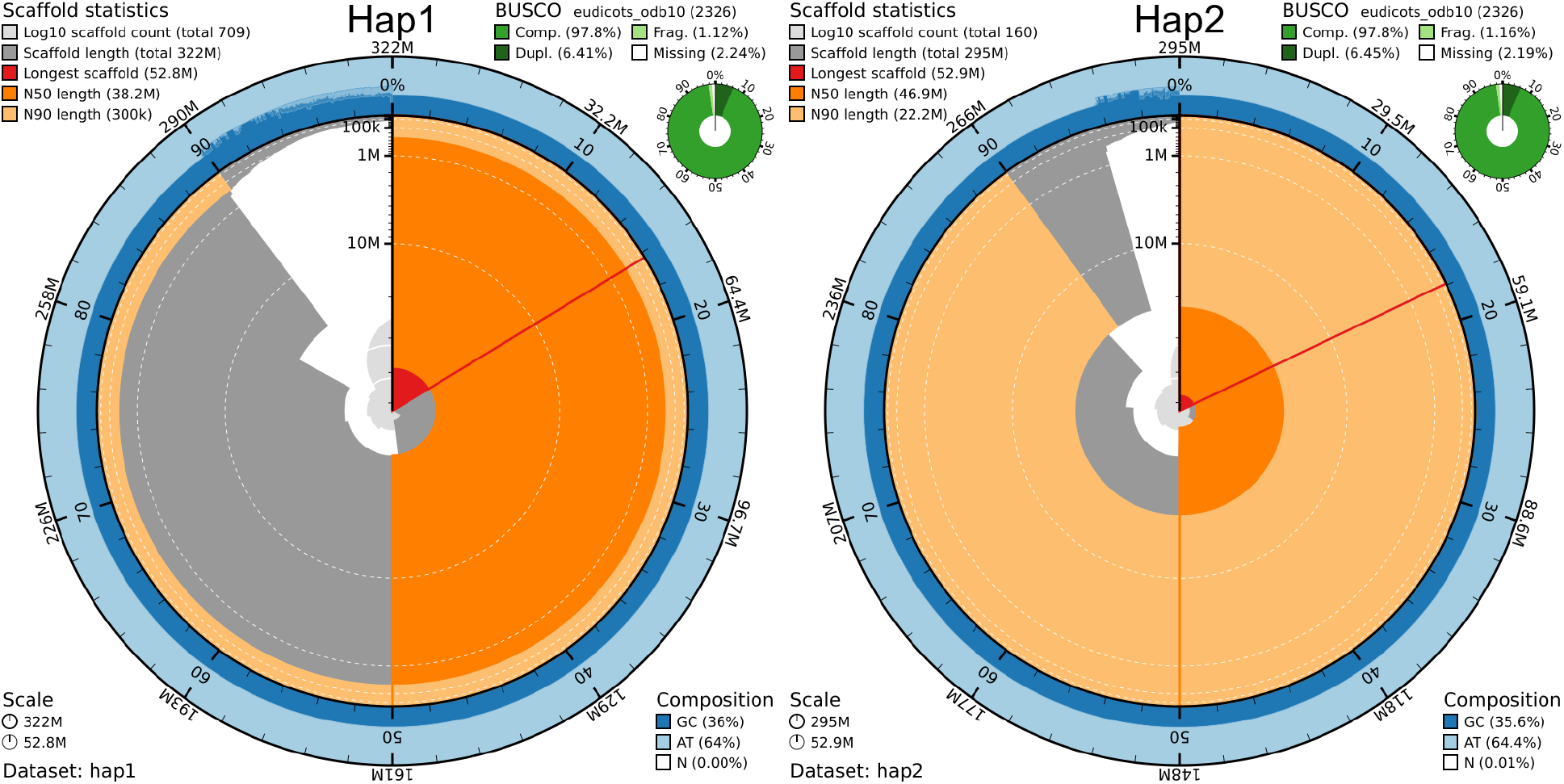
Metrics of the genome assemblies of *C. groenlandica* hap1 and hap2. BlobToolKit Snailplots show N50 metrics and BUSCO gene completeness. The two outermost bands of the circle signify GC versus AT composition at 0.1% intervals, with mean, maximum, and minimum. The third outermost shows the N90 scaffold length, while the fourth is N50 scaffold length. The red line shows the size of the largest scaffold. All the scaffolds are arranged in a clockwise manner from the largest to the smallest and are shown in darker gray with white lines at different orders of magnitude, while the light gray shows cumulative count of scaffolds.

When compared to a k-mer database of the Hi-C reads, hap1 had a k-mer completeness of 90.0%, hap2 of 89.4%, and combined they have a completeness of 90.6%. Further, hap1 had an assembly consensus quality value (QV) of 45.5 and hap2 of 51.3, where a QV of 40 corresponds to one error every 10,000 bp, or 99.99% accuracy compared to a k-mer database of the Hi-C reads (QV 56.6 and 61.6, respectively, compared to a k-mer database of the HiFi reads) (Table 2). A total of 34,213 and 29,684 protein-coding genes were annotated in hap1 and hap2, respectively (Table 2).

The Hi-C contact maps show clear separation of chromosomes into homologous sets (Supplementary Figure 3). The genome assemblies do not contain any detectable traces of contaminants (Supplementary Figure 4).

### Very large centromeric satellite arrays

Both assemblies contain extensive chromosomal regions enriched for tandem satellite repeats, accompanied by markedly reduced exon density (Figure 3, Supplementary Figure 5). These regions are consistent with exceptionally large centromeric or pericentromeric satellite arrays, although their precise functional boundaries remain undetermined, for example by CENH3 localization.

**Figure 3:**
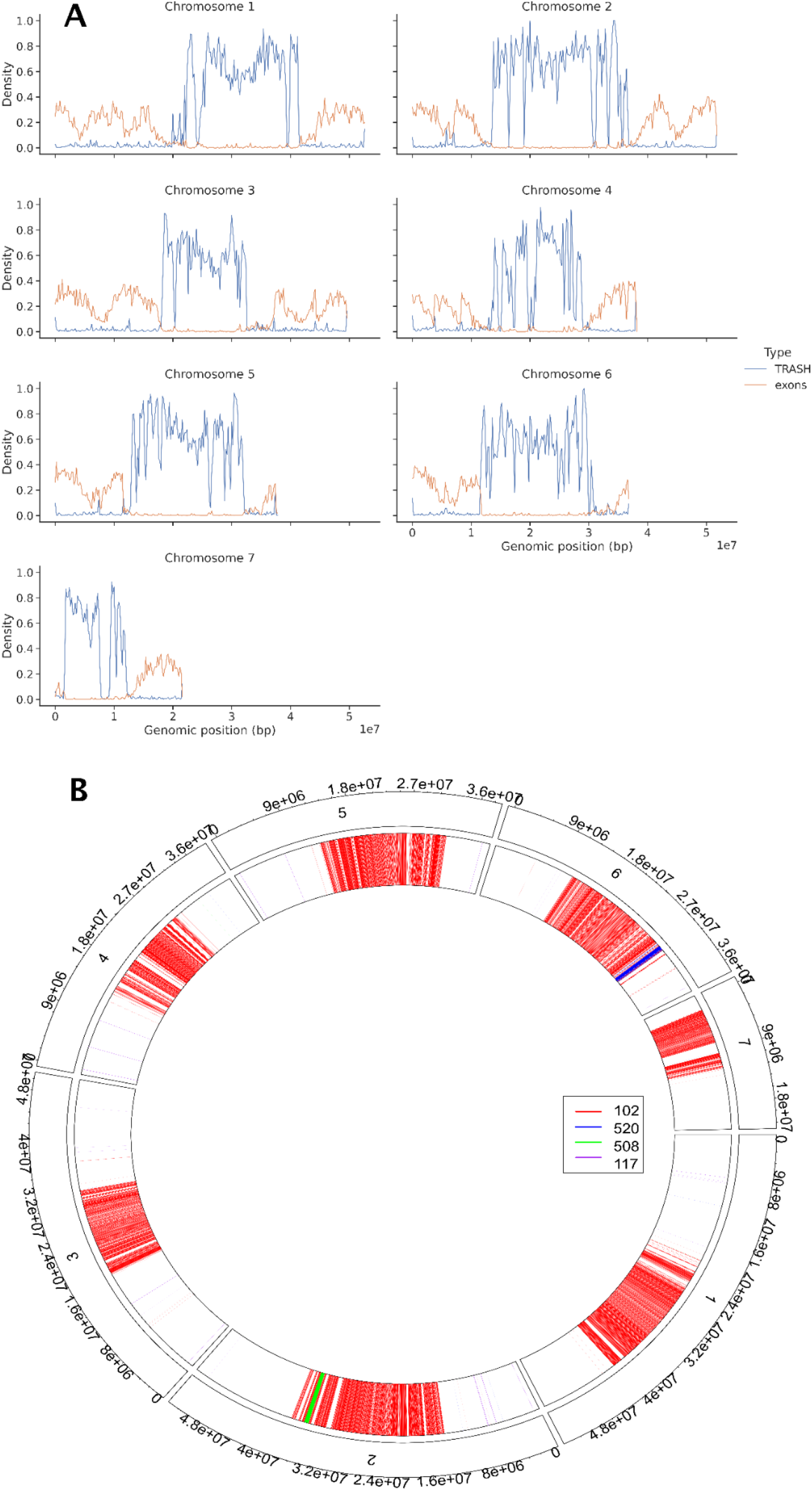
Extensive centromeric satellite arrays in *C. groenlandica*. A) Density of satellite sequences and exons along all seven chromosomes of hap1. Satellite sequences were identified with TRASH2 and exons densities were extracted from the gene annotation. B) Breakdown of repeat types consistent with centromeric arrays. Hap1 is presented as a circos plot. The tandem repeats found by TRASH are plotted in each chromosome. Supplementary Figure 5 shows the corresponding plots for hap2. Chromosome position is given on the outside of the circle and consensus repeat length is given in the internal rectangular key.

## Discussion

We have sequenced, assembled, and annotated chromosome-level, pseudo-haplotype separated genome assemblies of *C. groenlandica* originally collected from Alaska (as seeds) and grown in a growth chamber at University of Oslo. This is the first chromosome-level reference genome for a x=7 *Cochlearia* species and will be highly useful for population studies of this circumpolar distributed species. Furthermore, comparative analyses of chromosome evolution and structure of x=6 and x=7 (Gill, 1971; Saunte, 2010) diploid species as well as the various polyploids shaped by complex processes involving both allopolyploidy and autopolyploidy (Hämälä et al., 2024; M. Koch, 2002; Wolf et al., 2021) should be possible as soon as there are chromosome level assemblies for several additional the *Cochlearia* species.

A striking feature of the *C. groenlandica* genome is the exceptional extent of its putative centromeric satellite arrays, which occupy substantial portions of several chromosomes. Although these regions are consistent with unusually extensive centromeric or pericentromeric sequence, the exact location and extent of functional centromeres remain to be established Interestingly, previous work in *C. officinalis* identified strong signatures of selection associated with whole-genome duplication at kinetochore-associated genes, including *CENP-E, CENP-C, NDC80* and *INCENP* and additionally documented a very similarly vast preponderance of 102 bp putative centromeric satellite arrays in x=6 *Cochlearia excelsa* (Bray et al., 2024). Together, these observations suggest that centromere and kinetochore evolution may represent an interesting avenue for future investigation in the recurrently polyploid *Cochlearia*, for which the present assembly provides a valuable reference.

## Acknowledgments

This project received data management and infrastructure support from ELIXIR Norway, supported by the Research Council of Norway’s grants 322392 and 350529, the University of Bergen, the University of Oslo, the Arctic University of Norway in Tromsø, the Norwegian University of Science and Technology, and the Norwegian University of Life Sciences: NMBU. The authors acknowledge support from the National Infrastructure for High Performance Computing and resources provided by Sigma2 as well as Data Storage in Norway (project NN8013K) for computational work. The Norwegian Sequencing Centre generated the sequencing data used in this project (http://sequencing.uio.no).

## Data availability

Data generated for this study are available under ENA BioProject PRJEB90397. PacBio sequencing data for the Greenland scurvy-grass (ENA BioSample: SAMEA118445818) are deposited in ENA under ERR15104775, whereas Illumina Hi-C sequencing data are deposited in ENA under ERR15350460. Pseudo-haplotype one can be found in ENA at PRJEB90206, whereas hap2 is PRJEB90396.

The gene annotations are available at https://doi.org/10.5281/zenodo.22108523.

