## Supplementary information for "Chromosome-level genome assembly of Greenland scurvy-grass (*Cochlearia groenlandica*) reveals extensive centromeric satellite arrays"

Authors:
**Ole K. Tørresen [1], Anne Krag Brysting[1], Bram Danneels[2], Rhianah Sandean[3], Spyridon Kollias[1], Giada Ferrari [1], Ave Tooming-Klunderud [1], Levi Yant[3], Kjetill S. Jakobsen [1*]**
Addresses:
1) Department of Biosciences, University of Oslo, Oslo, Norway
2) Computational Biology Unit, Department of Informatics, University of Bergen, Bergen 5020, Norway
3) Linnean Centre for Plant Biology, Department of Plant Biology, Uppsala BioCenter, Swedish University of Agricultural Sciences, Uppsala, Sweden

### Supplementary information

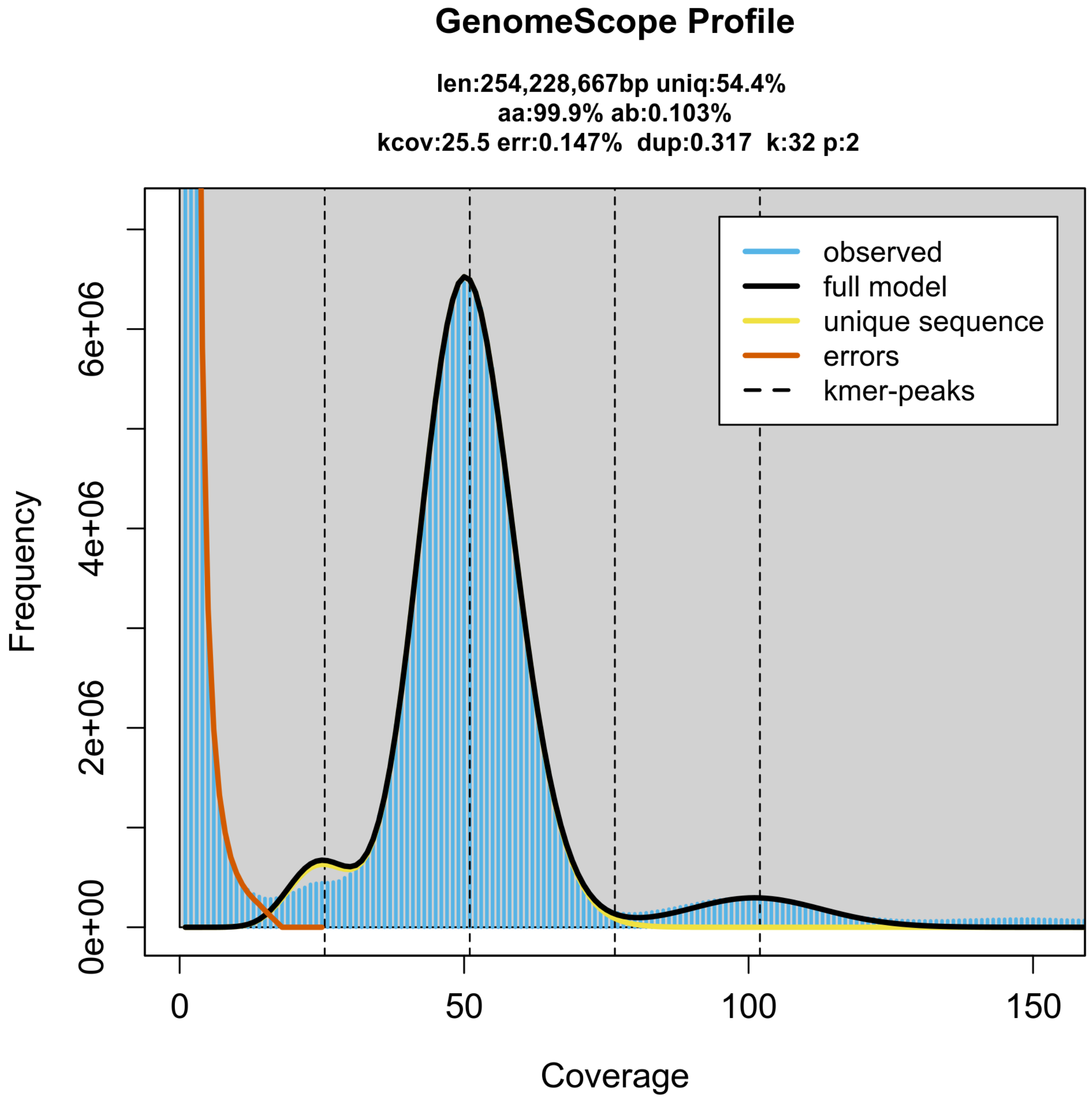


**Supplementary Figure 1: GenomeScope profile of the HiFi reads from the sequenced individual of *C. groenlandica*.** This analysis estimates a 254 Mb genome, with 0.1% heterozygosity.

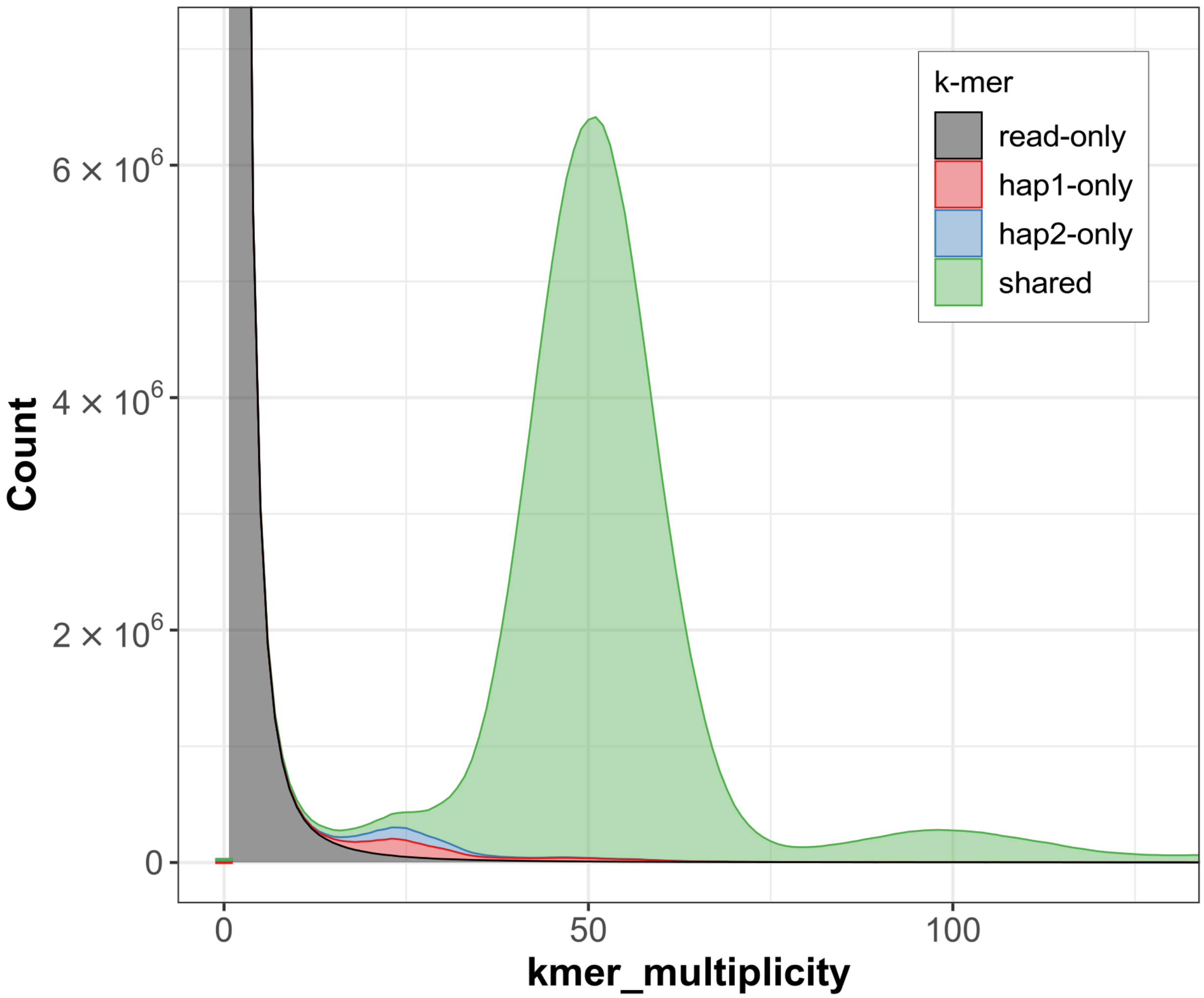


**Supplementary Figure 2: K-mer copy-number spectrum analysis of *C. groenlandica* compared to k-mers from a database from the HiFi reads.** Assembly-specific k-mers are shown in red and blue, while k-mers shared by both pseudo-haplotypes are in green. The stack above 0 on the x-axis shows k-mers found in the assemblies, but not in the reads. The figure is generated by Merqury.

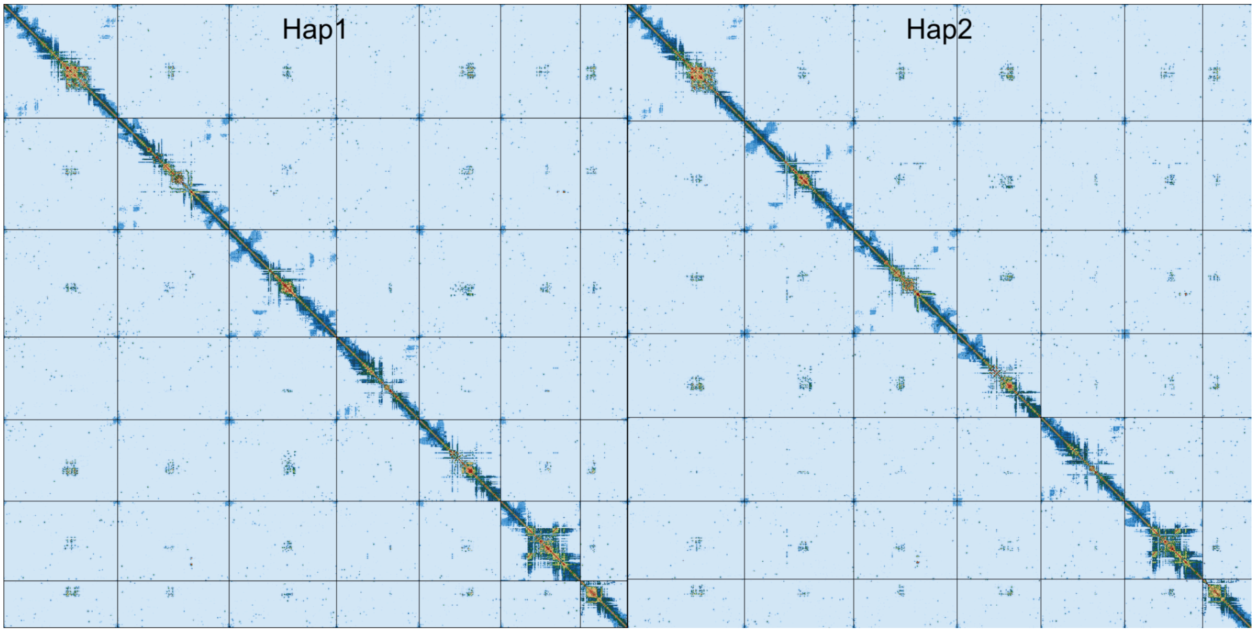


**Supplementary Figure 3. Hi-C contact map of genome assemblies of *C. groenlandica* for hap1 and hap2.** The assemblies are visualized using PreTextSnapshot. Chromosomes are shown in order of size from left to right and top to bottom.

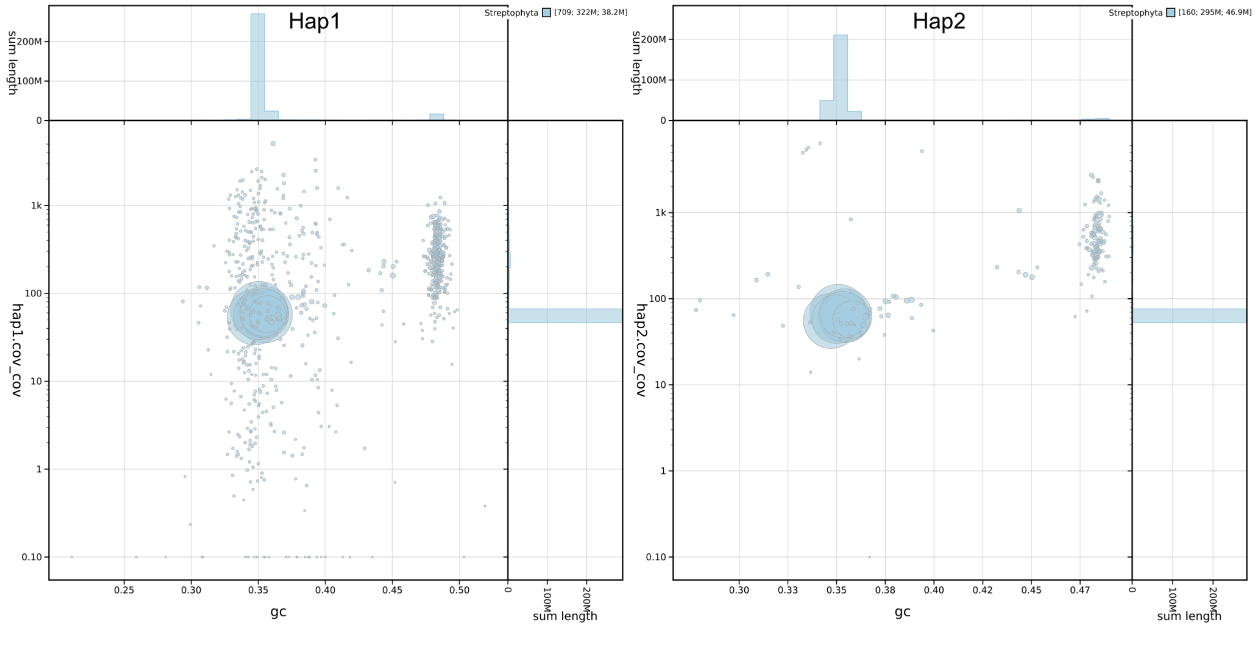


**Supplementary Figure 4: BlobToolKit GC-coverage plots of genome assemblies of *C. groenlandica* hap1 and hap2.** The scaffolds are coloured by phylum. The size of the circles are in proportion to the length of the scaffolds. Histograms show the distribution of scaffold length sum along each axis.

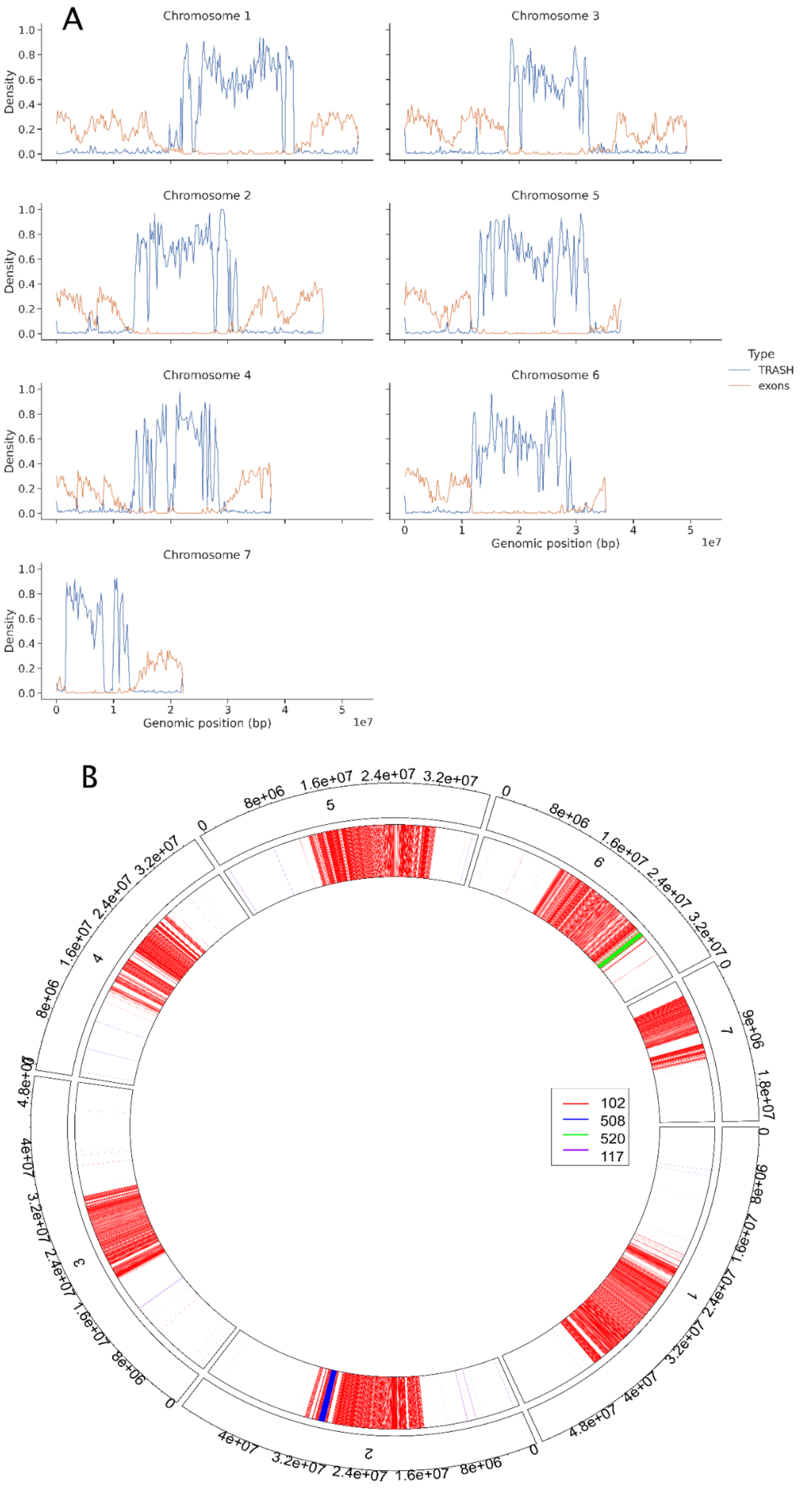
A
Chromosome 1
Chromosome 3
Chromosome 2
Chromosome 5
Chromosome 4
Chromosome 6
Chromosome 7
Density
1.0
0.8
0.6
0.4
0.2
0.0
0
1
2
3
4
5
1e7
Genomic position (bp)
Type
TRASH
exons
B
0
8e+06
1.6e+07
2.4e+07
3.2e+07
4e+07
4.8e+07
9e+06
1.8e+07
1
2
3
4
5
6
7
102
508
520
117

**Supplementary Figure 5: Extensive centromeric satellite arrays in *C. groenlandica*.** A) Density of satellite sequences and exons along all seven chromosomes of hap2. Satellite sequences were identified with TRASH2 and exons densities were extracted from the gene annotation. B) Breakdown of repeat types consistent with centromeric arrays. Hap2 is presented as a circos plot. The tandem repeats found by TRASH are plotted in each chromosome. Figure 3 shows the corresponding plots for hap1. Chromosome position is given on the outside of the circle and consensus repeat length is given in the internal rectangular key.
